# Chromosome-level genome assembly of the spider mite *Tetranychus cinnabarinus*

**DOI:** 10.1101/2025.06.24.661196

**Authors:** Kaiyang Feng, Xiang Wen, Lin He

## Abstract

*Tetranychus cinnabarinus* is an important agricultural pest, primarily due to its rapid development of pesticide resistance. In this study, we constructed a chromosome-level genome of *T. cinnabarinus* using Illumina, Nanopore, and Hi-C sequencing technologies. The assembled genome spans 86.67 Mb, with an N50 value of 21.77 Mb, and 84.54 Mb of sequences anchored across three chromosomes. The BUSCO completeness of the genome is 92.78%. Through gene prediction, we identified 11,388 protein-coding genes, 98.53% of which are functionally annotated. Furthermore, we systematically identified 304 genes from five major detoxification enzyme systems. This high-quality genome provides a solid foundation for investigating the physiological mechanisms of *T. cinnabarinus* and for identifying potential targets for biocontrol strategies.

## Background & Summary

*Tetranychus cinnabarinus* is an agricultural pest worldwide, infesting over 1,200 plant species(*1, 2*). Currently, control methods for this mite heavily depend on the application of chemical pesticides. However, the unique physiological traits of *T. cinnabarinus*, such as high reproductive capacity, significant generation overlap, and limited biological activity range combined with the unreasonable use of chemical pesticides have led to prominent resistance issues(*3*). According to the latest statistics from the Insecticide Resistance Action Committee (IRAC), phytophagous mites, including *T. cinnabarinus*, have developed resistance to 97 types of chemical pesticides, making them one of the most severely resistant arthropods(*4*). *T. cinnabarinus* and *Tetranychus urticae* share similarities in morphology, developmental stages, and genetic types(*5*). Despite the presence of reproductive isolation, some acarologists classify them as different color forms of the same species. Nevertheless, recent studies employing bioinformatics analysis have confirmed that they are sibling species(*6*).

The advancement of genomics has greatly expanded the scope and depth of study on physiological changes and pesticide-resistance mechanisms in arthropods(*7*). The genome of *T. urticae* is the first genomic dataset published for mite species. It was initially assembled using Sanger sequencing, resulting in a genome of 89.6 Mb with 640 scaffolds.(*8*) Furthermore, Wybouw et al. reassembled these Sanger sequences into three pseudo-chromosomes, further improving the genome’s utility(*9*). In 2024, the genome of T. urticae was updated once again, with Cao et al. successfully generating a chromosome-level genome database through third-generation sequencing and Hi-C technology(*10*). This genome database has been extensively utilized in study on mite resistance, host adaptation, horizontal gene transfer, and growth and development, underscoring the crucial role of genomics in acarology. In this study, we successfully assembled a chromosome-level genome of *T. cinnabarinus* using Nanopore long-read sequencing, Illumina short-read sequencing, and Hi-C technology. The assembled genome is 86.69 Mb in size, with a contig N50 of 21.77 Mb. This high-quality genome provides valuable genetic resources for study on various mite species.

## Methods

### Mites strain

The original strain of *T. cinnabarinus* used in this study was collected from Beibei, Chongqing, China, in 1995(*11*). To minimize heterozygosity, a homozygous line with a same background was established. Briefly, an unfertilized female mite was selected for parthenogenetic reproduction to generate F1 male mites, which were then backcrossed with the maternal females to produce the F2 generation. In the F2 generation, an unfertilized female mite was again selected for parthenogenetic reproduction and backcrossing. This process of genotype purification was carried out for 15 generations, ultimately yielding a homozygous line with a same background.

### DNA extraction and sequencing

200 mg of female adult mites were selected, frozen in liquid nitrogen, and then genomic DNA was extracted using the QIAamp DNA Blood Mini Kit, following the manufacturer’s instructions. 2 μg of genomic DNA (gDNA) were repaired using the NEB Next FFPE DNA Repair Mix kit (M6630, USA) and then processed with the ONT Template Prep Kit (SQK-LSK109, UK) according to the manufacturer’s instructions. The resulting large segment library was premixed with loading beads and loaded into a previously used and washed R9 flow cell. Sequencing was performed on the ONT PromethION platform using the corresponding R9 cell and ONT sequencing reagent kit (EXP-FLP001.PRO.6, UK), following the manufacturer’s guidelines. The Hi-C library was sequenced on the NovaSeq 6000 platform. Briefly, adapter sequences from the raw reads were trimmed, and low-quality paired-end (PE) reads were removed to generate clean data. The resulting Hi-C clean reads, totaling 11.35 Gb (∼130×), were first truncated at the putative Hi-C junctions. The trimmed reads were then aligned to the assembly using the BWA aligner(*12, 13*). Only uniquely mappable paired reads with a mapping quality greater than 20 were retained for further analysis. Invalid read pairs, including dangling ends, self-cycles, re-ligation products, and dumped products, were filtered out using HiC-Pro v2.8.1(*14*). Final pseudo-chromosomes were constructed after manual adjustments, with the vast majority (97.54%) of the assembled sequences anchored to the three pseudo-chromosomes, 99.60% of which were ordered and oriented.

### Genome survey

A genome survey was conducted using a *k-mer*-based method(*15*). The *k-mer* coverage was determined from Illumina short reads using Jellyfish version 2.2.10 with a *k-mer* size of 19(*15*). Genome size, heterozygosity, and duplication rate were estimated using GenomeScope version 2.0(*16*). The estimated genome size of *T. cinnabarinus* is 97.29 Mb, with a heterozygosity rate of 0.05% (Fig. 1). The proportion of repetitive sequences is approximately 26.30%, while the GC content is about 33.04%.

**Fig. 1.**
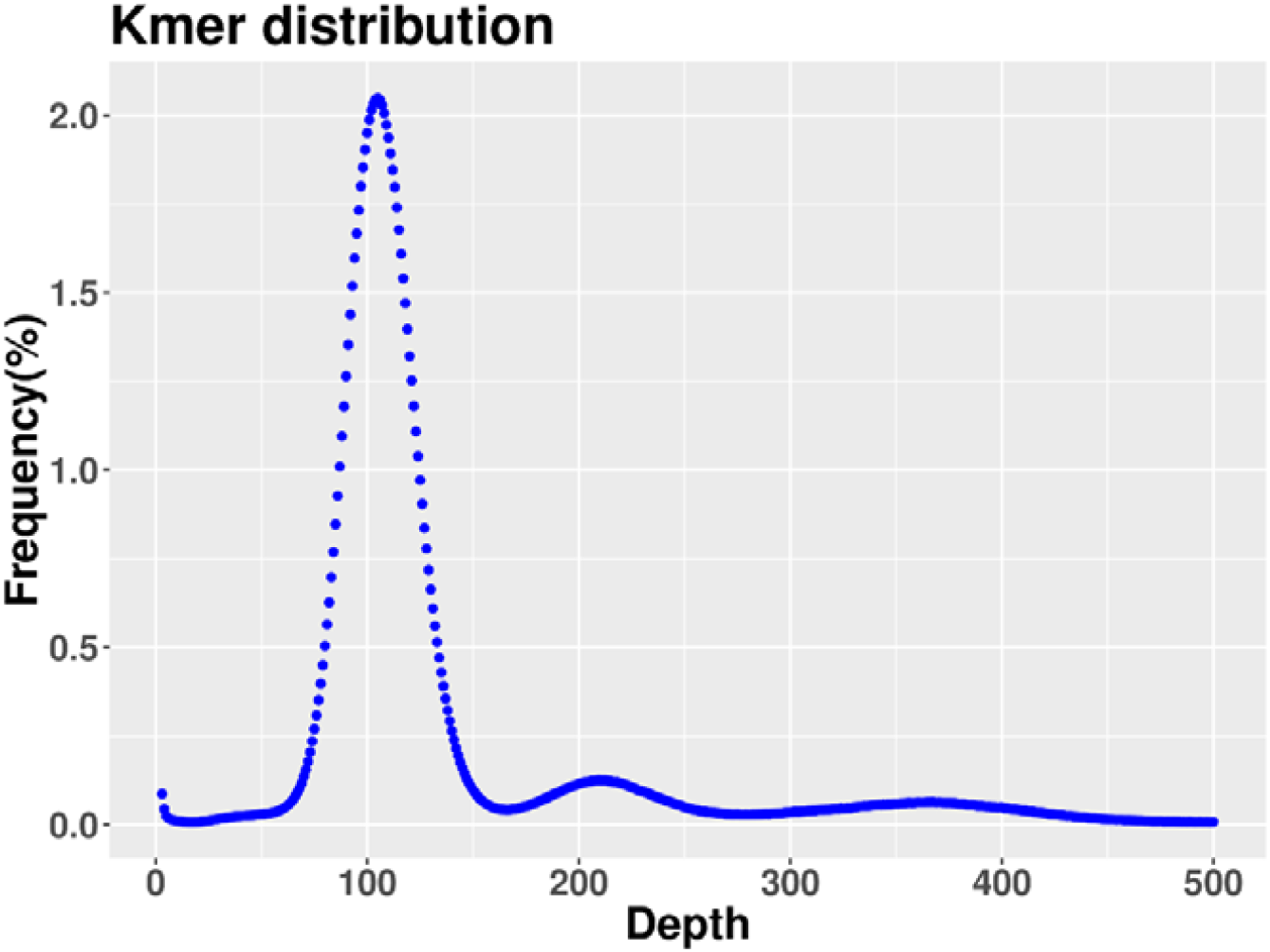
Genome survey of *T. cinnabarinus* Genome size, heterozygosity and rate of duplication were estimated using Genomescope. *k-mer□*= □19.

### Genome assembly

In this study, NeCat software (https://github.com/xiaochuanle/NECAT) was employed for the error correction and assembly of third-generation Nanopore sequencing data(*17*). Subsequently, PURGE_DUPS software was used to eliminate contigs with high heterozygosity(*18*). NeCat software was then used for iterative correction of the preliminary assembly results using second-generation Illumina data, enhancing the overall assembly quality across two rounds. To generate high-quality genomic resources, Hi-C technology was also employed for assisted assembly. Using Juicer software, which interfaces with the BWA program, Hi-C reads were aligned to the initial genomic draft sequence, followed by filtering with a restriction endonuclease (Dpn II) tag. The 3D-DNA software, operated with default parameters, facilitated the error correction, anchoring, ordering, and locating of contig fragments, resulting in chromosome-level assembly outcomes(*19*). Following the output from 3D-DNA, the assembly results were visualized using Juicebox (v1.11.08), with manual adjustments made including correction of chromosome boundaries, erroneous connections, and translocations and inversions(*20*). The quality of the genome was assessed using BUSCO (Benchmarking Universal Single Copy Orthologs) software(*21*). A total of 1,066 BUSCOs from the Insecta_odb10 dataset were aligned against the assembled genome, employing multiple indicators based on a hidden Markov model to evaluate genome completeness, including complete and single-copy, complete and duplicated, fragmented, and missing sequences.

After de novo assembly, 39 contigs (N50 of 21.77□Mb) with a total length of 86.67□Mb were generated (Table 1). The Hi-C chromosome mapping results indicate that a total of 84.54 Mb of sequences were assigned to three chromosomes, representing 97.54% of the total. Of the sequences mapped to the chromosomes, 84.20 Mb were successfully ordered and oriented, constituting 99.60% of the total mapped sequence length (Table 1 and Fig. 2). The GC content of the *T. cinnabarinus* genome was 32.34% (Table 1), which is similar to that of *T. urticae* (32.25%) and *T. piercei* (32.17%)(*22*).

**Table 1.**
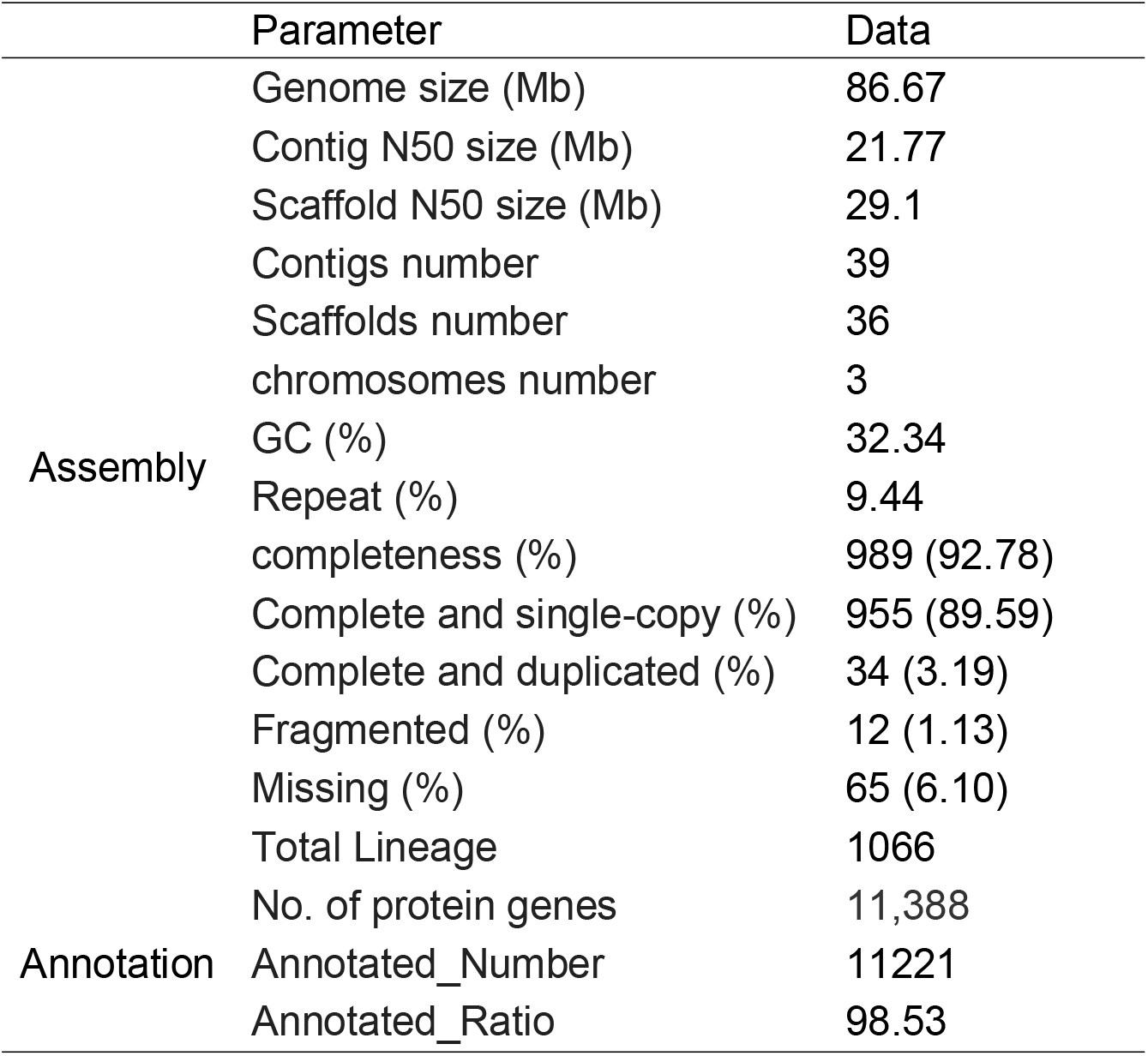
Genome assembly and annotation information of *T. cinnabarinus*.

|  | Parameter | Data |
| --- | --- | --- |
| Assembly | Genome size (Mb) | 86.67 |
|  | Contig N50 size (Mb) | 21.77 |
|  | Scaffold N50 size (Mb) | 29.1 |
|  | Contigs number | 39 |
|  | Scaffolds number | 36 |
|  | chromosomes number | 3 |
|  | GC (%) | 32.34 |
|  | Repeat (%) | 9.44 |
|  | completeness (%) | 989 (92.78) |
|  | Complete and single-copy (%) | 955 (89.59) |
|  | Complete and duplicated (%) | 34 (3.19) |
|  | Fragmented (%) | 12 (1.13) |
|  | Missing (%) | 65 (6.10) |
|  | Total Lineage | 1066 |
| Annotation | No. of protein genes | 11,388 |
|  | Annotated_Number | 11221 |
|  | Annotated_Ratio | 98.53 |

**Fig. 2.**
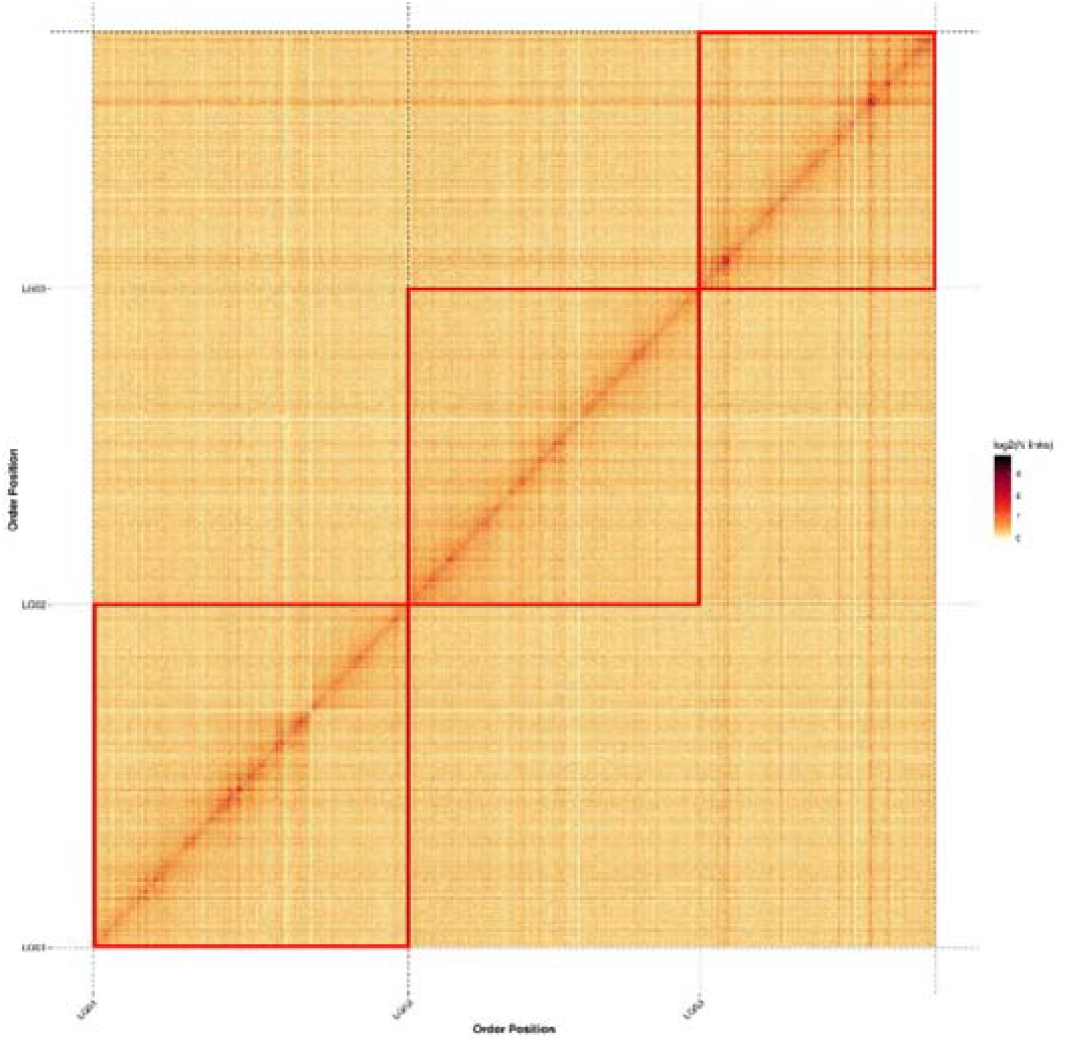
Genome-wide chromosomal heatmap of *T. cinnabarinus* The red boxes indicate three linkage groups.

### Genome annotation

A de novo repeat library for *T. cinnabarinus* was constructed using RepeatModeler v2.0.2a(*23*), focusing on the specific structures of repetitive sequences. This custom library was further refined by integrating data from Dfam 3.3 and RepBase-20181026(*24, 25*). Repetitive sequences were subsequently identified through searches conducted with RepeatMasker v4.1.2p(*26*). RNA-seq data were used to aid in the prediction of protein-coding genes. The RNA-seq reads were mapped to *T. cinnabarinus* genome assembly using Hisat v2.2.0(*27*), followed by transcript assembly using Stringtie v2.1.2(*28*). For genome annotation, gene prediction was conducted using SNAP v2013-02-16 and Augustus v3.2.3(*29, 30*), with the resulting parameters further trained and refined. The predicted protein-coding genes were compared with genomic data from *Drosophila melanogaster* and *T. urticae* for homologous analysis(*7, 8*), leading to the creation of a protein-coding gene database. Functional annotations of the predicted gene sequences were subsequently performed using the NR, EggNOG, GO, KEGG, SWISS-PROT, and Pfam databases. In this study, we predicted 11,388 protein-coding genes from *T. cinnabarinus* genome (Table 1), with 98.53% of these genes annotated in the databases.

### Identification of detoxification enzyme genes

Previous studies have shown that the amplification of detoxification enzyme genes is a key factor contributing to the strong host adaptability and rapid development of pesticide resistance in spider mites(*31*). Therefore, this study further investigates the detoxification enzyme genes in *T. cinnabarinus* genome(*32-36*). In this study, we identified a total of 282 genes from five major detoxification enzyme systems, including 72 cytochrome P450 (P450) genes (Fig. 3a), 25 glutathione S-transferases (GST) genes (Fig. 3b), 57 carboxylesterase (CCE) genes (Fig. 3c), 44 UDP-glucuronosyltransferase (UGT) genes (Fig. 3d), and 84 ABC transporter (ABC) genes (Fig. 3e).

**Fig. 3.**
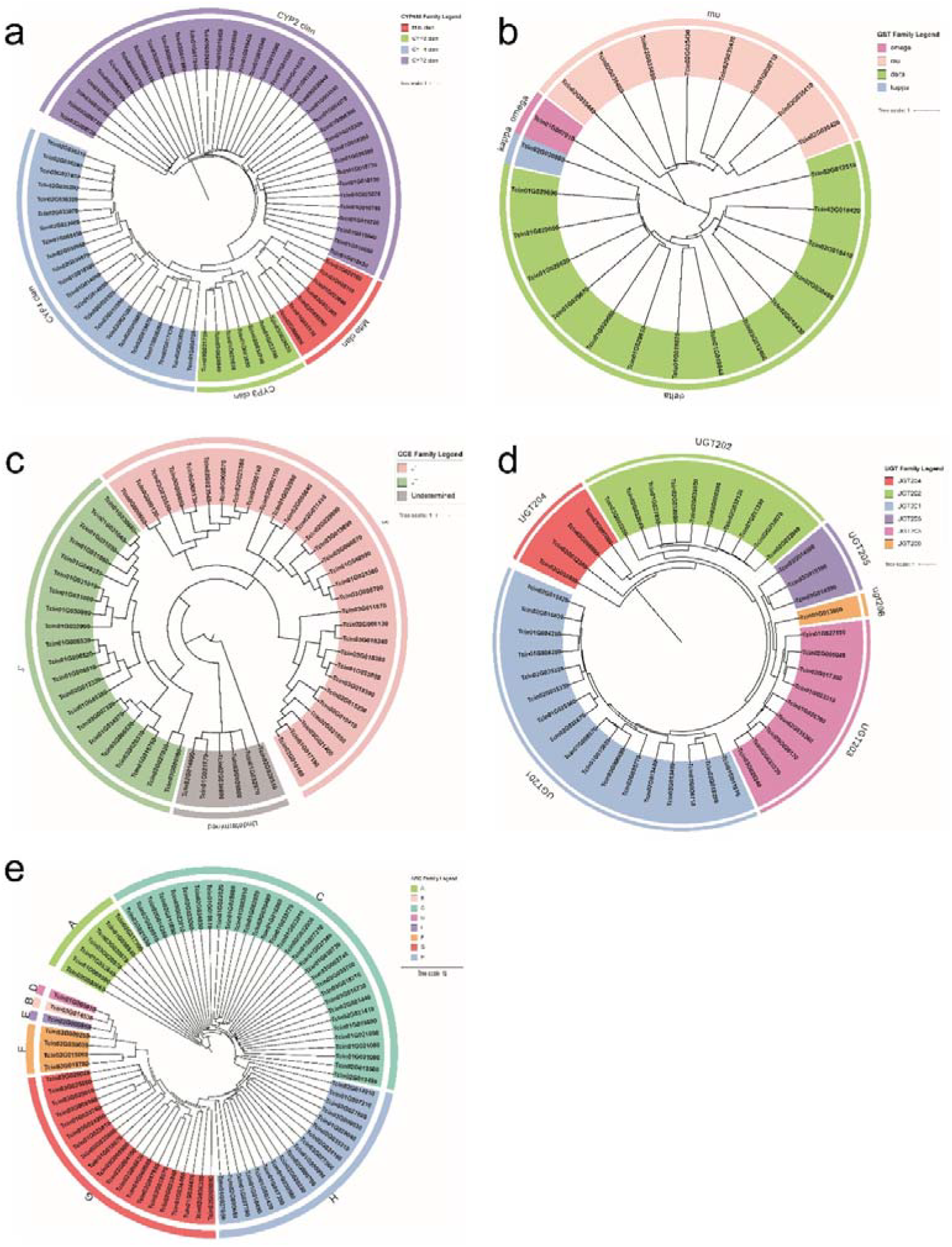
The detoxification enzyme genes in *T. cinnabarinus* genome (a) P450 genes phylogenetic analysis, (b) GST genes phylogenetic analysis, (c) CCE genes phylogenetic analysis, (d) UGT genes phylogenetic analysis, (e) ABC genes phylogenetic analysis.

### Data Records

This Whole Genome Shotgun project has been deposited at GenBank under the accession JBLVUC000000000. The *T. cinnabarinus* genome project has been submitted to NCBI under BioProject No. PRJNA1227279. The version described in this paper is version JBLVUC010000000.

### Technical Validation

The completeness of the genome was evaluated using 1066 conserved core genes from the OrthoDB v11 database(*37*). The results indicated that *T. cinnabarinus* genome achieved a completeness of 92.78% (989), with 89.59% (955) classified as complete and single-copy, 3.19% (34) as complete and duplicated, 1.13% (12) as fragmented, and 6.10% (65) as missing sequences (Table 1). This genome completeness is comparable to that of the published *T. urticae* and *T. piercei* genome(*9, 10*). Furthermore, mapping of RNA-seq Illumina short reads to the genome revealed a mapping rate of 97.11%, which further confirms the high completeness of *T. cinnabarinus* genome. In conclusion, we have successfully assembled a high-quality chromosome-level genome for *T. cinnabarinus*.

## Acknowledgements

This study was supported by grants from the National Natural Science Foundation of China (32202337 and U2202202), the Fundamental Research Funds for the Central Universities (SWU-KQ23018).

